# Rapid value learning reveals generalized and context-dependent codes in frontal cortex

**DOI:** 10.64898/2026.07.03.736027

**Authors:** Elena Gutierrez, Timothy H. Muller, James L. Butler, W. M. Nishantha Malalasekera, Laurence T. Hunt, Sebastijan Veselic, Steven W. Kennerley

## Abstract

Studying how the brain represents value spans distinct methods and training histories, from neuroimaging in task-naïve humans to single-neuron recordings in extensively trained non-human primates. Similar findings across fields have encouraged the untested assumption that rapidly emerging and overtrained value representations are equivalent. Here we recorded single-neuron activity in anterior cingulate cortex (ACC) and orbitofrontal cortex (OFC) as macaques learned novel cue values and chose between novel and overtrained cues. Value responses emerged within 4-7 cue presentations, matching behavioural adaptation. Yet ACC and OFC used distinct codes. ACC encoded value in a common format that generalised across training history. OFC coding was more context-dependent, with distinct subpopulations recruited by choice experience. Choice novelty was also encoded independently of value before chosen value signals emerged. During learning, ACC responses shifted from overtrained secondary reinforcers to newly predictive cues. These findings establish the necessary (rapid acquisition) and sufficient (generalised format) conditions for comparing value representations across methods, species, and training histories.

## Introduction

The ability to assign value to newly encountered options is essential to flexible choice. The anterior cingulate (ACC) and orbitofrontal cortex (OFC) have been consistently implicated in value-guided computations in both human and non-human primates (Ballesta et al., 2020; Enel et al., 2020; Hunt et al., 2018; Kennerley et al., 2011; Klein-Flugge et al., 2022; Lee et al., 2012). While studies in both species largely support each other’s findings, methodologies are at odds. In non-human primates (NHPs), single-cell data have been mostly obtained once animals reach a high-performance criterion after experiencing the task, choices and task conditions thousands of times. More recent studies have examined changes in neural responses as animals acquire novel value-guided tasks from scratch, but still typically do so across weeks of extensive training (Liebana et al., 2025; Wojcik et al., 2026). Though extensive training ensures precise behavioural control, this bears little resemblance to human neuroimaging experiments, where participants often experience the task for the first time on the day of data collection. As a result, it remains a prominent but largely untested assumption that the overtrained representations of value reported in NHP neurons underlie the newly-learned representations of value observed in human neuroimaging studies. Demonstrating such similarity would therefore not only strengthen evidence for cross-method and cross-species parallelisms, but also reveal the type of rapid value acquisition that is essential to adaptive decision-making.

The assumption that novel and overtrained value representations are similar is non-trivial, because value-related responses in frontal cortex (FC) are modulated by novelty, experience, and task state (Barron et al., 2013; Costa & Averbeck, 2020; Elston & Wallis, 2025; Wilson et al., 2014), raising the possibility that newly acquired and overtrained values rely on distinct neural codes. If, for instance, objects are tagged as novel, increased attentional resources may be devoted to them to reduce uncertainty in the environment, thus maximising future reward. Indeed, incorporating novelty in reinforcement learning algorithms has proven effective in driving learning through increased exploration (Jaegle et al., 2019). The FC has been shown to be involved in novelty detection through interaction with the medial temporal lobe (Daffner et al., 2000; Garrido et al., 2015; Kafkas & Montaldi, 2014; Matsumoto, Matsumoto, & Tanaka, 2007; Ranganath & Rainer, 2003). Neurons in the OFC particularly support exploration of novel options (Costa & Averbeck, 2020) and, in humans, the medial frontal cortex is preferentially recruited during consideration of novel experiences (Barron et al., 2013), with value-related signals here diminishing with training (Hunt et al., 2012). Taken together, these results demonstrate novelty may be a salient signal for PFC neurons. More broadly, the fact that FC responses are modulated by experience (Asaad et al., 1998; Cole et al., 2010; Rainer & Miller, 2000), and that OFC responses may be state-dependent (Elston & Wallis, 2025; Wilson et al., 2014), reinforces the question of whether novelty and training experience also affect value-guided computations. Consistent with this, Bongioanni et al. (2021) used functional MRI to show how population responses in the medial frontal cortex may encode the value of novel options by presenting NHPs with previously unencountered combinations of reward-predicting stimuli. Extending a similar approach to electrophysiological recordings may therefore reveal how neurons encode novelty, whether these representations are independent of value-related signals and/or whether novel choice differentially biases value responses of frontal subregions, shedding light on the neural mechanisms that support rapid, flexible choice.

More fundamentally, if novelty and training experience shape frontal value coding, how are value representations acquired as novel stimuli become behaviourally meaningful? One influential framework for value learning, temporal-difference reinforcement learning (TDRL), predicts that as reward becomes associated with a cue, value-related neural responses should shift backwards in time from the reward itself to earlier reward-predicting events (Amo et al., 2022; Sutton, 1988). Mechanisms consistent with this account have been extensively characterised in midbrain dopamine neurons, which encode reward prediction errors (RPEs) by signalling the discrepancy between predicted and obtained reward (Schultz et al., 1993; Schultz et al., 1997), and can acquire responses to novel reward-predicting stimuli over repeated exposures via secondary reinforcers (Hollerman & Schultz, 1998). Because the midbrain dopamine system projects widely to frontal cortex (Williams & Goldman-Rakic, 1998), and particularly to ACC (Averbeck et al., 2014), similar error-driven dynamics may contribute to the acquisition of value representations in frontal neurons. This possibility is supported by evidence that ACC disruption impairs the integration of reinforcement information during learning (Amiez et al., 2006; Camille et al., 2011; Kennerley et al., 2006), that ACC neurons encode RPEs during action- and rule-learning (Cole et al., 2024; Matsumoto, Matsumoto, et al., 2007; Seo & Lee, 2007), and that ACC value responses reflect discrepancies between expected and obtained reward during choice (Kennerley et al., 2011; Muller et al., 2024). Crucially, however, these findings largely involve cues, actions or task structures that had already been extensively experienced before recordings began. Therefore, while our task does not by itself identify the precise learning algorithm, it allows us to test whether a core signature associated with TDRL (i.e. the temporal transfer of value coding from known reward-predicting events to newly learned cues) is evident in frontal neurons, and whether such dynamics occur over timescales fast enough to support rapid behavioural adaptation.

Here, we addressed these issues by recording the activity of single neurons in ACC and OFC while two non-human primates (*Macaca mulatta*) first learned novel cue-value associations through secondary conditioning, and then made choices between the newly learned cues and/or overtrained cues. We report the clear emergence of value signals associated with the novel cues as rapidly as within just four cue presentations, on approximately the same timescale as behavioural adaptation. In ACC neurons, the association of value with novel cues coincided with a decrease in value coding at the secondary reinforcer, as predicted by TDRL. Upon choice, ACC used a strikingly similar value code for both novel and overtrained choices, whereas OFC recruited distinct subpopulations according to level of experience. OFC also strongly encoded novelty of the choice independently of value.

## Results

Two subjects (F and M) learned novel stimuli associated with five different sizes or probabilities of reward (Fig. 1a-c). Each session (F: *n* = 8, M: *n* = 13), ten novel cues were learned through completion of 100 Conditioning trials (‘Conditioning phase’, Fig. 1a-b), consisting of forced choice of a cue, followed by presentation of a secondary reinforcer which represented the reward magnitude or probability predicted by the cue. Conditioning trials were grouped into ten blocks for each attribute, consisting of five Conditioning trials – one for each reward level – as well as two Probe Choice trials between cues from that attribute (*n* = 40/session). Subjects were rapidly able to choose the correct option, performing above chance within 3-5 cue presentations (mean accuracy ± S.E.M. on Probe Choice trials, averaged across sessions, F: 70.3 ± 5.7% by 5-6 presentations, *p* = 0.007, M: 63.5 ± 5.2% by 3-4 presentations, *p* = 0.041; one-sample *t*-tests, Bonferroni-corrected within subject; Fig. 2b). A notable feature of the secondary reinforcer is that it allows for equally rapid acquisition of both probability and magnitude information. Multiple trials are typically required to reliably estimate probability information (but not magnitude); the presence of the secondary reinforcer removes this ambiguity, as the height of the secondary reinforcer reflects the true expected value of the primary reinforcer. Consistent with this, Probe choice performance did not significantly differ between magnitude and probability trials during either early or late conditioning (F, presentations 1-3: *t*(7) = −2.35, *p* = 0.10, presentations 7-10: *t*(7) = 0, *p* = 1; M, presentations 1-3: *t*(12) = 0.50, *p* = 1, presentations 7-10: *t*(12) = 2.51, *p* = 0.06; paired *t*-test, Bonferroni-corrected within-subject; Fig. 2c); we therefore collapsed across these two types of stimuli in many of the subsequent analyses.

**Figure 1.**
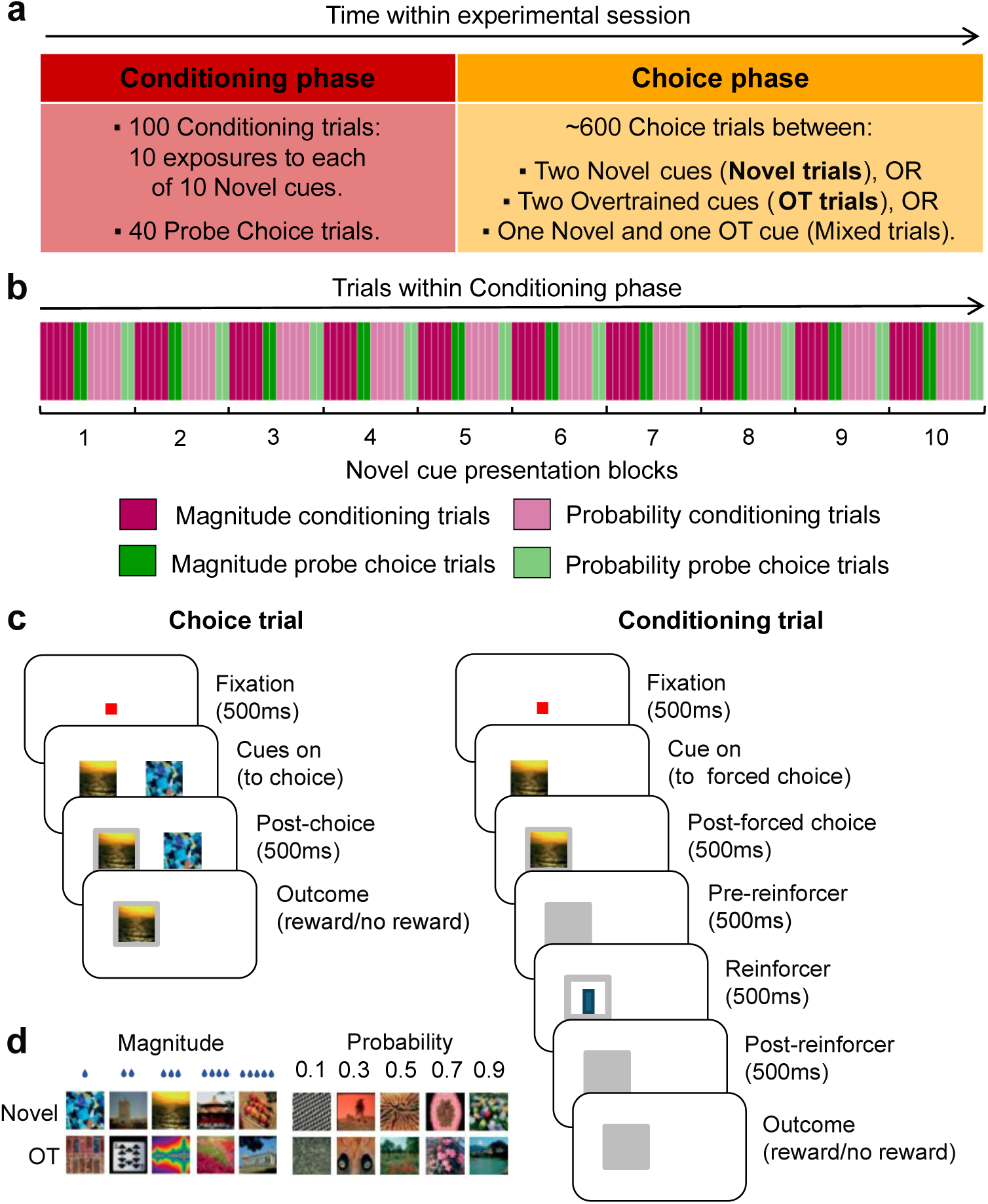
Task. **a**, Task design. During each session, subjects learnt the value associated with novel cues predicting different sizes or probabilities of reward (‘Conditioning phase’). Subjects were then offered choices between two cues they learned that session (Novel trials), between two cues they experienced 1000s of times before (Overtrained trials) or between one of each (Mixed trials) (‘Choice phase’). **b**, Conditioning phase structure. Each novel cue was presented 10 times, for a total of 10 blocks for each attribute (magnitude and probability). Each block consisted of a Conditioning trial for each of the 5 stimuli of an attribute, followed by two probe Choice trials from this attribute. Blocks alternated between magnitude and probability trials, with the first being randomly determined (magnitude in this example). **c**, Left: in Choice trials, subjects chose between two cues of the same attribute (magnitude or probability) by joystick movement, followed by reward outcome associated with the chosen cue. Right: during Conditioning trials, each novel cue was paired with an overlearned secondary reinforcer which indicated the novel cue’s value. Corresponding reward outcome followed a forced choice of the cue by manual joystick movement. **d**, Example stimuli.

**Figure 2.**
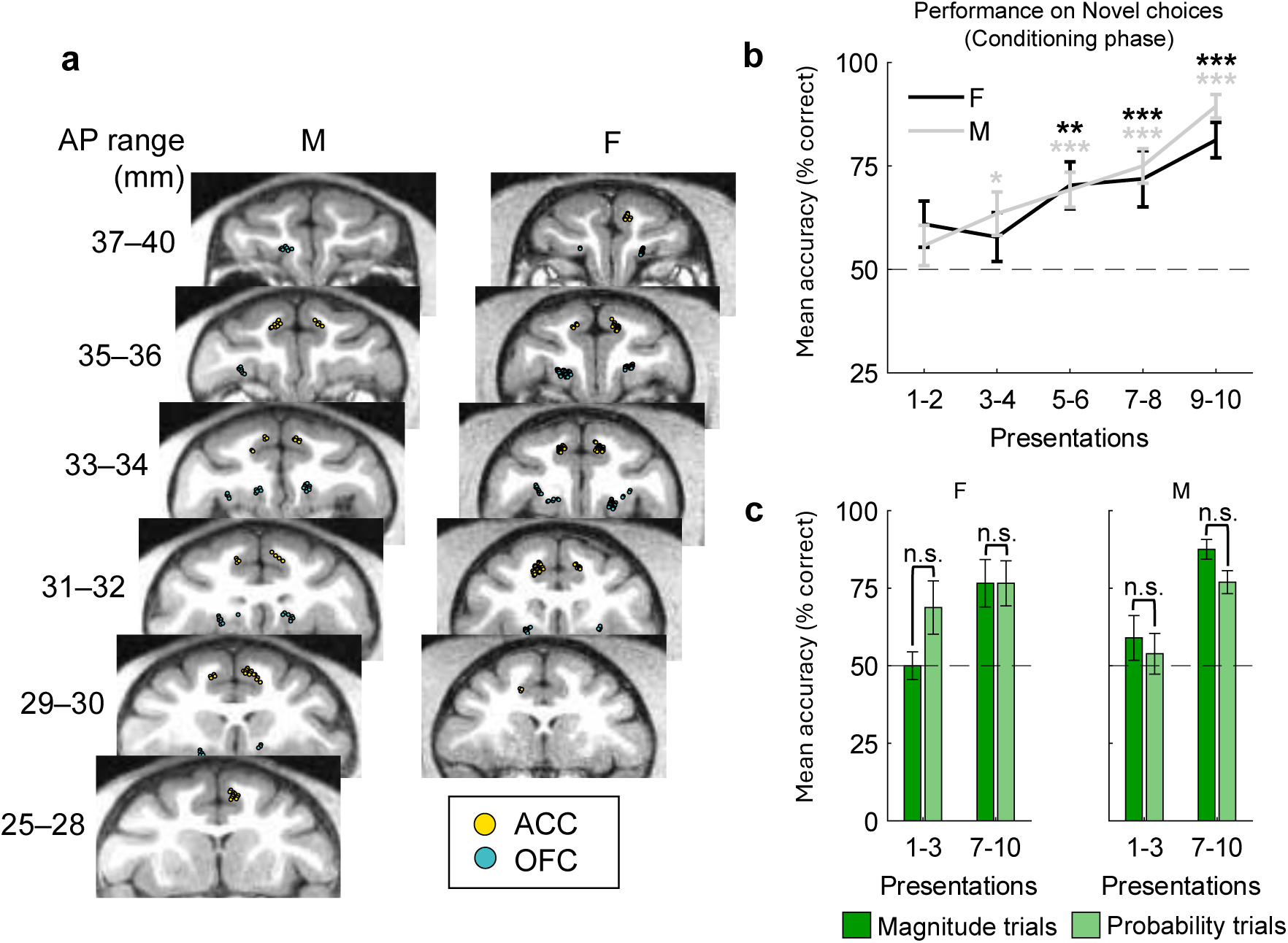
Recording locations and learning behaviour. **a**, Recording locations of ACC and OFC neurons, shown in coronal sections. Anterior–posterior (AP) range indicates the position anterior to the interaural plane in stereotactic coordinates. **b-c**, Conditioning phase choice behaviour on novel stimuli (Probe Choice trials). **b**, Improvement in accuracy across presentation blocks (grouped in five bins of 8 Probe Choice trials, or 2 presentation blocks, each) averaged across sessions (±SEM). \**p*<0.05, \*\**p*<0.01, \*\*\**p*<0.0001, one-sample *t*-test, Bonferroni-corrected. **c**, Accuracy in the early (presentations 1-3) and late (presentations 7-10) Probe Choice trials of the Conditioning phase, averaged across sessions (±SEM). Trials were split according to attribute.

The Conditioning phase was followed by a Choice phase, where subjects were presented with novel choices (between cues learned that day, Novel trials) or overtrained choices (between the same, extensively learned cues, Overtrained, or OT, trials). After cue presentation, subjects could freely view the stimuli until they chose to indicate their decision with a lever pull (mean reaction time ± SEM, F: 547 ± 7ms, M: 481 ± 2ms). We previously reported on behavioural analyses of the Choice phase, finding that initial saccades during choice were value-driven, and that NHPs have a strong bias towards fixating novel options even in the absence of any bias towards choosing them (Cavanagh et al., 2019). Here, we report neuronal activity from 156 neurons located in ACC (primarily from the dorsal bank of the anterior cingulate sulcus, area 24c) and 160 neurons located in OFC (primarily from the medial orbital gyrus, area 11/13) (Fig. 2a). We focus on whether the choice context (novel vs overtrained) impacts value-related signals, and how value-related signals emerge during learning.

### Overtraining does not alter ACC value representations, but recruits distinct OFC subpopulations

Presenting choices between the cues learned that day (Novel trials) or between heavily experienced cues (OT trials) allowed us to evaluate whether training alters the nature of ACC and OFC value representations. Note that for each of the five levels of value, the reward probability (or reward magnitude) associated with a Novel or OT stimulus was identical; only the amount of experience with that stimulus differed (Fig. 1a).

We first examined how strongly neurons encoded chosen value in Novel and OT trials (GLM-1). Value signals emerged in ACC and OFC ∼200ms post cue-onset and showed no significant difference between Novel and OT trials (coefficient of partial determination, CPD; permutation test; Fig. 3a, Supplementary Fig. 1). In both regions, selectivity strength increased across trial epochs (ACC: *F*(1, 155) = 19.87, *p* < 0.0001, OFC: *F*(1, 157) = 6.58, *p* = 0.011; repeated-measures ANOVA), whereas choice experience had no effect on value coding, indicating that value explained a comparable fraction of firing rate variance across experience levels. Consistent with this, choice experience did not produce a systematic change in population firing rate (mean FR ± SEM, averaged across neurons and epochs, ACC: 12.14 ± 1.05 on Novel trials and 12.14 ± 1.04 on OT trials, *t*(155) = −0.01, *p* = 0.9; OFC: 9.41 ± 0.74 on Novel trials and 9.32 ± 0.74 on OT trials, *t*(159) = 0.75, *p* = 0.5).

**Figure 3.**
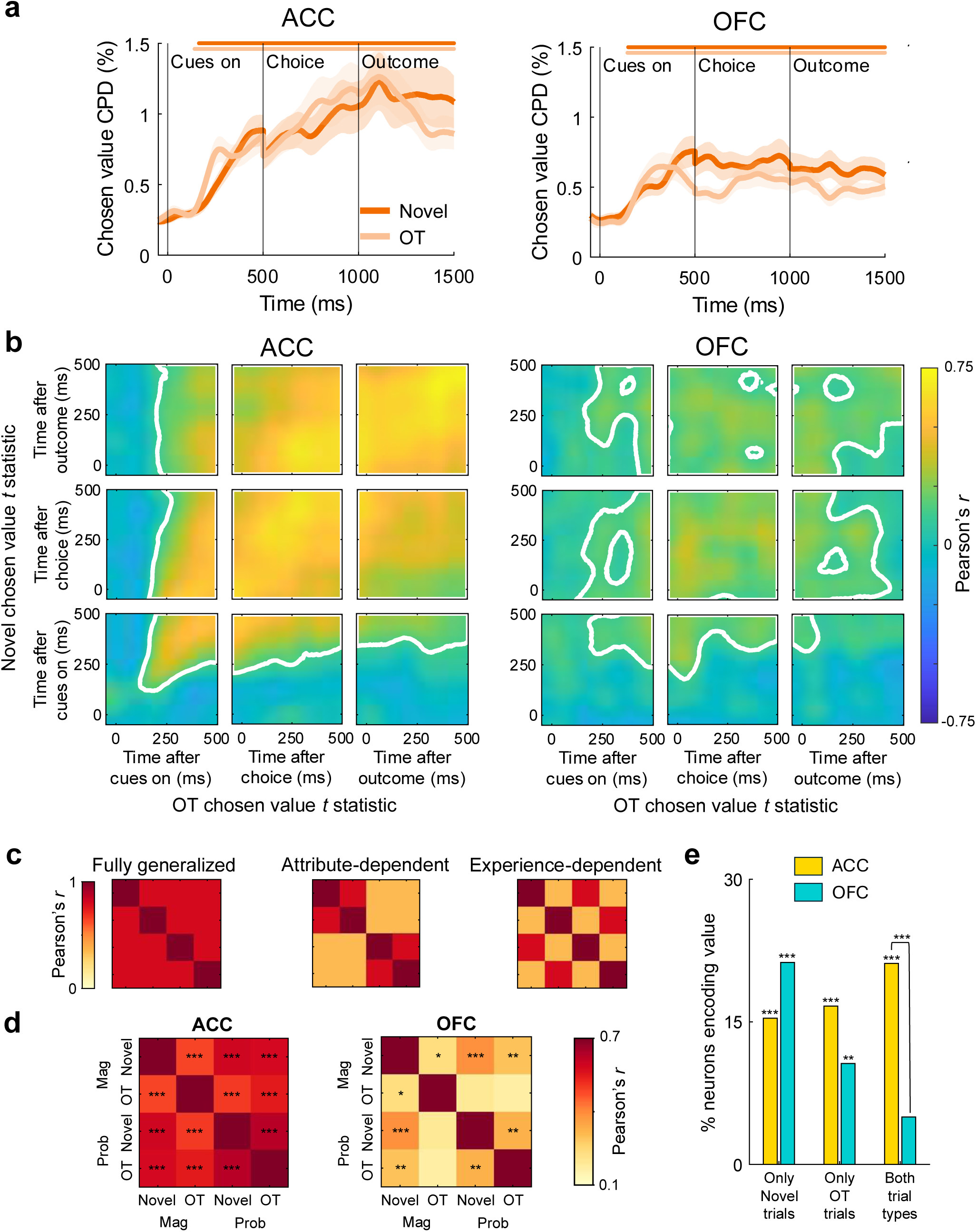
Choice experience does not alter ACC value representations but recruits distinct OFC subpopulations. **a**, Coefficient of partial determination (CPD, %) for chosen value during the Choice phase, split between Novel and OT choice trials, averaged across all ACC (left) and OFC (right) neurons. Coloured horizontal lines denote significance within the corresponding trial split, determined by permutation testing. **b**, Cross-correlation matrices reflecting the time-varying relationship between value coding (chosen value *t* statistics) in Novel and OT trials across all ACC (left) and OFC (right) neurons. White lines denote significance determined by cluster-based permutation testing. **c**, Template correlation matrices illustrating candidate organizations of the chosen value code across reward attribute and choice experience: fully generalized (top), attribute-dependent (or generalizing over experience, middle), and experience-dependent (or generalizing over attribute, bottom). Conditions are ordered as Novel magnitude, OT magnitude, Novel probability, and OT probability. **d**, Empirical correlation matrices for value coding in ACC (left) and OFC (right). Each cell shows the Pearson correlation between neuron-wise chosen value *t* statistics for the corresponding pair of conditions, estimated from firing rates averaged across the 1.5 s following cue onset across all neurons within each region. ACC exhibits a more fully generalized value code than OFC, whereas OFC shows evidence of an attribute-dependent code specifically during OT trials. \**p*<0.05, \*\**p*<0.001, \*\*\**p*<0.0001, Bonferroni-corrected. **e**, Proportion of ACC and OFC neurons encoding value only in Novel trials, only in OT trials or in both trial types. \**p*<0.05, \*\**p*<0.001, \*\*\**p*<0.0001, Binomial test and Chi-squared test.

To further explore whether value representations were sensitive to choice novelty, we examined the cross-correlation of *t* statistics for chosen value on Novel and OT trials (GLM-1). This revealed a strikingly positive correlation in the ACC population, which emerged approximately at the same time as value signals (∼200ms post-cues on) and was stably maintained for the entire trial duration (cluster-based permutation test; Fig. 3b), suggesting the value representations used in both Novel and OT choices were similar. While Novel and OT value coding was also positively correlated in OFC (Fig. 3b), this correlation was significantly lower than in ACC (cluster-based permutation tests; Supplementary Fig. 2). Thus, while both regions encoded the value of Novel and OT choices equally strongly, ACC neurons used a generalised value code across Novel and OT cues when compared to OFC neurons.

Having found stronger Novel-OT generalisation in ACC than OFC, we next asked how this experience-related structure relates to reward attribute. ACC and OFC are known to differ in how their value codes generalise across choice attributes, with ACC tending to support a more generalised code and OFC more often showing context- and attribute-specific coding (Elston & Wallis, 2025; Kennerley et al., 2009). We therefore summarised value coding across the same post-cue choice epoch shown in Fig. 3b, and examined chosen value correlations across the four trial types defined by reward attribute (magnitude or probability) and experience (Novel or OT). We considered three possible organisations of the value code (Fig. 3c): a ‘fully generalised’ code, predicting similar value coding across all conditions; an ‘attribute-dependent’ code, predicting shared value coding between Novel and OT trials within the same attribute; or an ‘experience-dependent’ code, predicting shared value coding between magnitude and probability trials within the same level of experience. ACC more closely resembled a fully generalised code than OFC. In ACC, chosen value representations showed significant correlations both across experience and across attribute, consistent with a common-currency format that generalised across both task dimensions (Fig. 3d; Supplementary Fig. 3). In OFC, by contrast, value representations remained more segregated across task dimensions: correlations remained significantly lower than those in ACC for all comparisons, and magnitude-probability correlations within OT trials were not significant, consistent with attribute-dependent value coding. These findings suggest that ACC generalises value coding across both reward attribute and choice experience, whereas OFC retains a more structured code consistent with context- and attribute-specificity, though this pattern only emerges with experience.

What is the source of OFC’s experience-dependent value code? Examination of value-selective neurons (GLM-1; Fig. 3e) revealed OFC neurons tended to be selective either in Novel or OT trials, rather than both, consistent with a more experience-dependent code. ACC neurons were in fact more likely to encode value across both trial types (*χ*² = 18.25, *p* < 0.0001; Supplementary Fig. 4). Overall, this pattern is further evidence that ACC maintains a more generalised value code across experience, whereas OFC relies on more distinct, experience-dependent subpopulations.

### Choice experience is encoded independently of value

Given the experience-dependent value code evident in OFC, we next asked whether ACC and OFC encoded experience with a choice context beyond its value. Such novelty signals could mark a choice context as unfamiliar, supporting the recruitment of attentional resources or promoting exploration under uncertainty. We thus tested whether choice novelty *per se* (i.e. independent of value) was encoded in ACC and OFC neurons during Novel and OT trials (GLM-1).

Both regions strongly encoded whether the choice offered was between Novel or between OT cues (Fig. 4a; Supplementary Fig. 5), particularly OFC, whose neurons were more likely to encode choice novelty than ACC neurons both before and after choice (choice: *χ^2^* = 12.26, *p* = 0.0009, outcome: *χ^2^* = 5.07, *p* = 0.048, Bonferroni-corrected; Fig. 4b). Choice experience is thus a salient contextual variable that modulates frontal neurons independently of chosen value (Fig. 4c).

**Figure 4.**
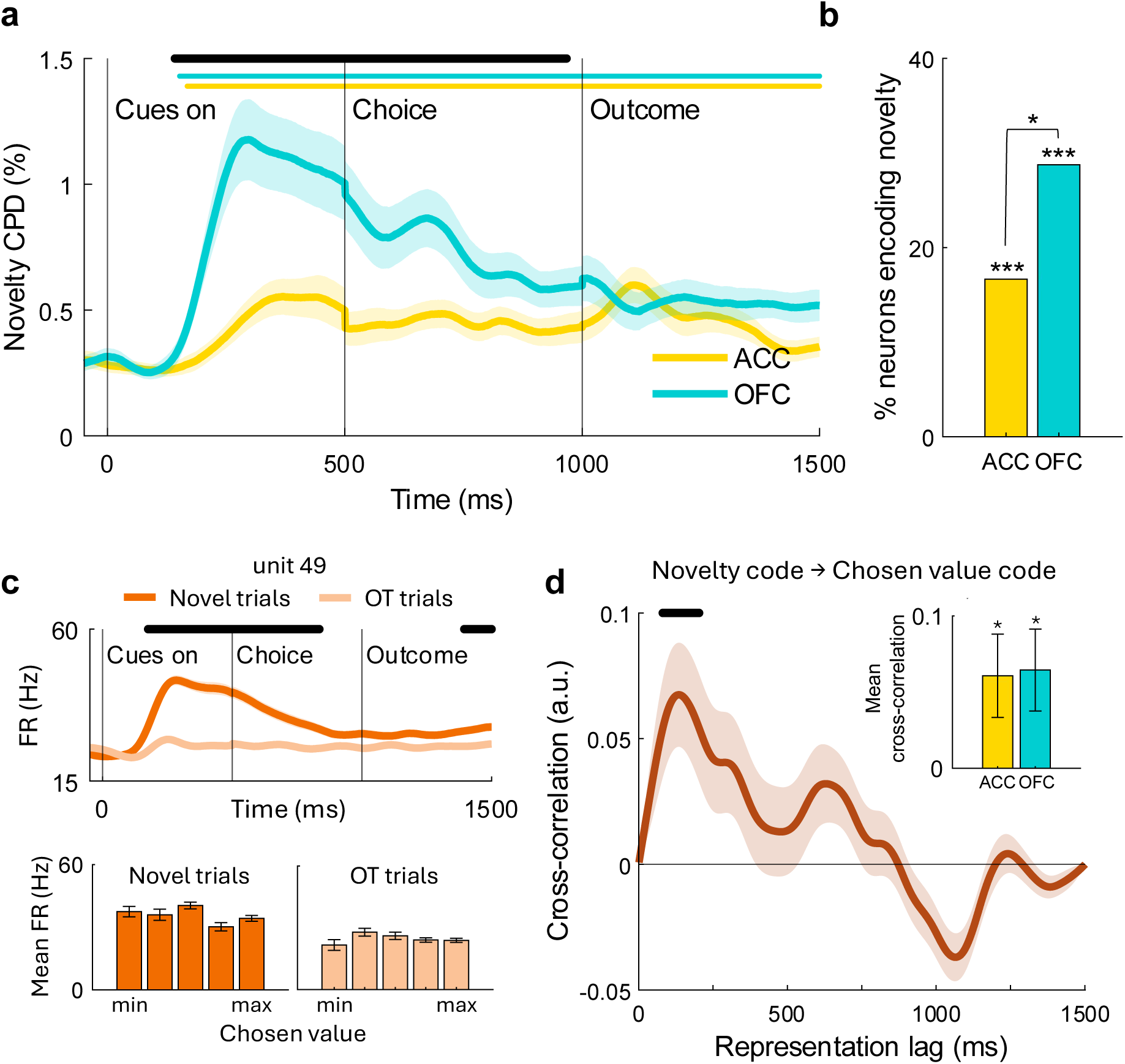
OFC encodes choice novelty independently of value. **a**, CPD (%) for choice novelty during novel and overtrained trials averaged across all ACC and OFC neurons. Colored horizontal lines denote significance within the corresponding region, black horizontal line denotes significant difference between regions, determined by permutation testing. **b**, Proportion of ACC and OFC neurons encoding novelty across trial epochs. \**p<*0.05, \*\*\**p*<0.001, chi squared test, Bonferroni-corrected; all *p*<0.0001 for binomial tests. **c**, Spike density function of an example OFC unit showing a difference in activity as a function of choice novelty. Solid horizontal line represents significant novelty coding (|*t*-stat|≥3.1 for >20ms). Bottom row: firing rate for the unit in top row averaged 100-500ms post-cue presentation in Novel and OT trials split according to the value of the chosen cue, showing no value-guided modulation of activity. **d**, Cross-correlation between the time courses of the novelty and chosen value signals averaged across all ACC and OFC neurons. Values greater than zero indicate stronger alignment when novelty coding preceded chosen value coding. Shading denotes SEM, black horizontal line indicates significant lags determined by permutation testing. Inset: mean cross-correlation averaged across the significant lags, shown separately for ACC and OFC neurons. Error bars denote SEM. \**p* < 0.05, one-sample *t* test.

If the choice novelty code is used to provide contextual input to support the recruitment of additional attentional resources, this signal ought to precede chosen value coding in time. To test this, we cross-correlated novelty and chosen value CPD time courses during Novel trials and quantified their temporal asymmetry (see Methods). Consistent with the prediction, choice novelty coding reliably preceded chosen value coding, with the cross-correlation peaking at a lag of ∼130 ms (Fig. 4d). Across both regions, novelty information thus emerged before value signals, consistent with choice experience providing a contextual scaffold for subsequent value computation.

### Novel value representations emerge rapidly with learning

The introduction of novel cues in every session further allowed us to investigate the learning dynamics of value associations in ACC and OFC neurons. We first asked how rapidly value representations emerge in single neurons, in order to test whether they may arise on timescales similar to those observed in typical human neuroimaging experiments. We split Conditioning trials into three bins with a sliding window, equivalent to cue presentation blocks 1-4, 4-7 and 7-10 of each cue (*n* = 40 trials/bin; GLM-2; Fig. 1a-c). We focussed on comparison between the cue presentation and the secondary reinforcer epoch, as the latter were stimuli associated with the same value as the preceding novel cue, but which had been extensively experienced ahead of recordings. Value signals emerged post-cue within 4-7 presentations of the novel stimuli in ACC and within 7-10 presentations in OFC (permutation tests; Fig. 5). Signals arose ∼250ms after cue presentation in both regions, a timing consistent with current option evaluation (Hunt et al., 2018; Kennerley et al., 2011). Value post-secondary reinforcer and outcome, on the other hand, was already encoded within the first 1-4 presentations, reflecting the fact these stimuli were heavily overtrained. This was consistent across attributes (Supplementary Fig. 6). As rapidly as within 4-7 cue presentations, the activity of frontal neurons thus reflected the emergence of value representations associated with novel cues.

**Figure 5.**
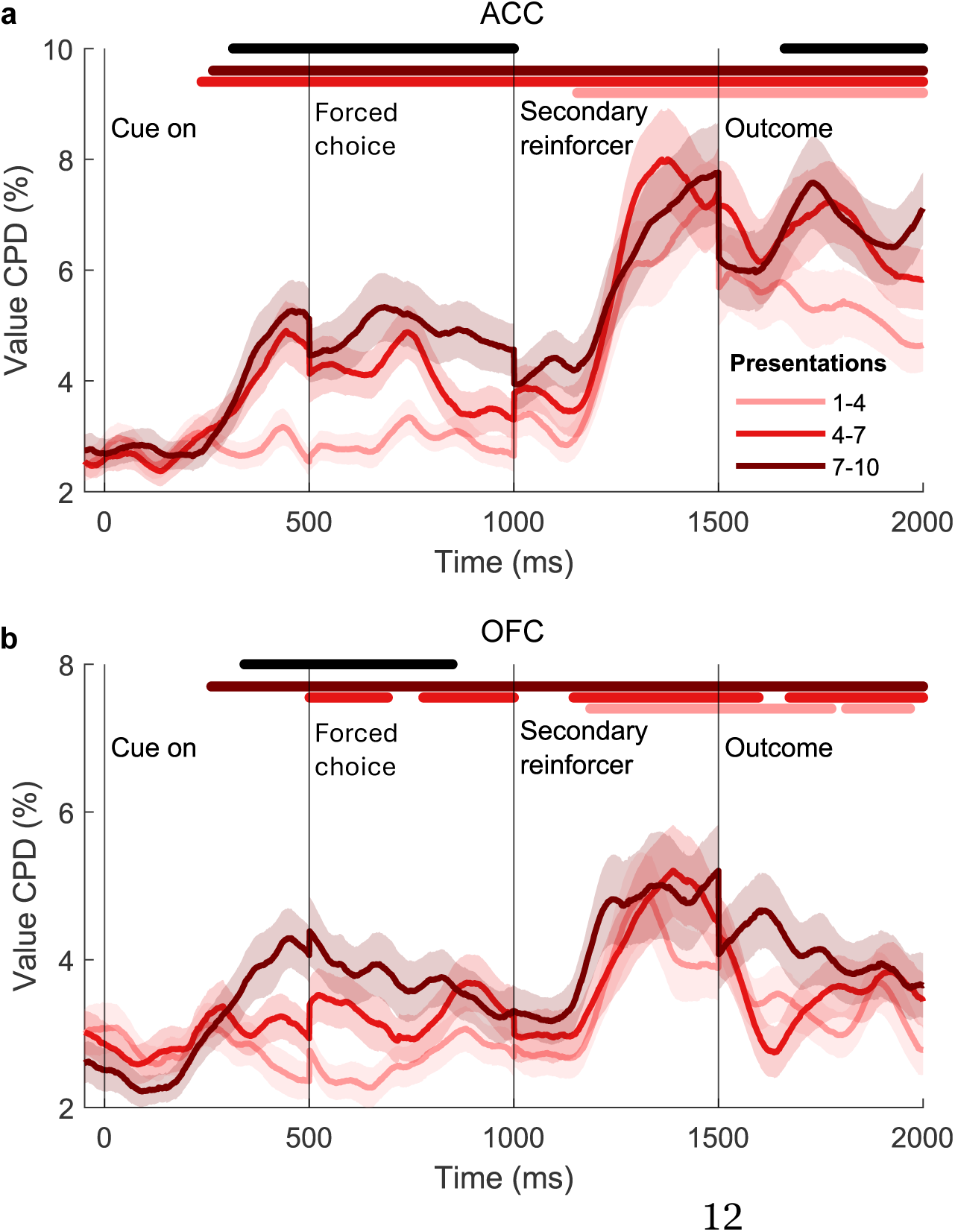
Novel value representations emerge rapidly with learning. **a-b**, CPD (%) for cue value during Conditioning trials, averaged across all ACC (**a**) and OFC (**b**) neurons. Trials were split in three bins comprising four cue presentations each using a sliding window (n=40 trials/bin). Horizontal lines denote significance determined by permutation testing. Coloured lines denote significance within trial-split, black horizontal line denotes a significant difference between presentations 1-4 and 7-10.

### ACC neurons demonstrate RPE-driven learning dynamics

We next asked how value representations were acquired. A hallmark of reward prediction error (RPE)-based learning is the backpropagation of value signals from conditioned stimuli or rewards, onto earlier stimuli or states that predict the reward (Sutton & Barto, 1987). Within single neurons, this may manifest in the firing rate initially correlating with value only after presentation of the overlearned reward-predicting stimuli; with some experience of the association of the novel cue with reward, the value signal should then emerge at the presentation of the newly learned cue (Fig. 6a). We explored whether frontal neurons exhibited signatures of RPE-based learning by examining Conditioning trials, where precise information about the value associated with a novel cue could be extracted following presentation of the secondary reinforcer. To test whether the emergent value representations at cue resembled those at secondary reinforcer, we compared value coding at cue during presentations 7-10 (‘Late’) to value coding at secondary reinforcer during presentations 1-4 (‘Early’; GLM-2). Cross-correlating Late value *t* statistics at cue to Early value *t* statistics at secondary reinforcer revealed a positive correlation between Late cue and Early secondary reinforcer value coding in both ACC and OFC (permutation tests, Fig. 6b). This indicated that, as learning progressed, value coding at the newly-learned cue came to reflect the pattern previously observed at the secondary reinforcer. As predicted, the opposite relationship – Early value coding at cue to Late value coding at secondary reinforcer – showed no correlation, likely due to the absence of cue-value coding in early trials.

**Figure 6.**
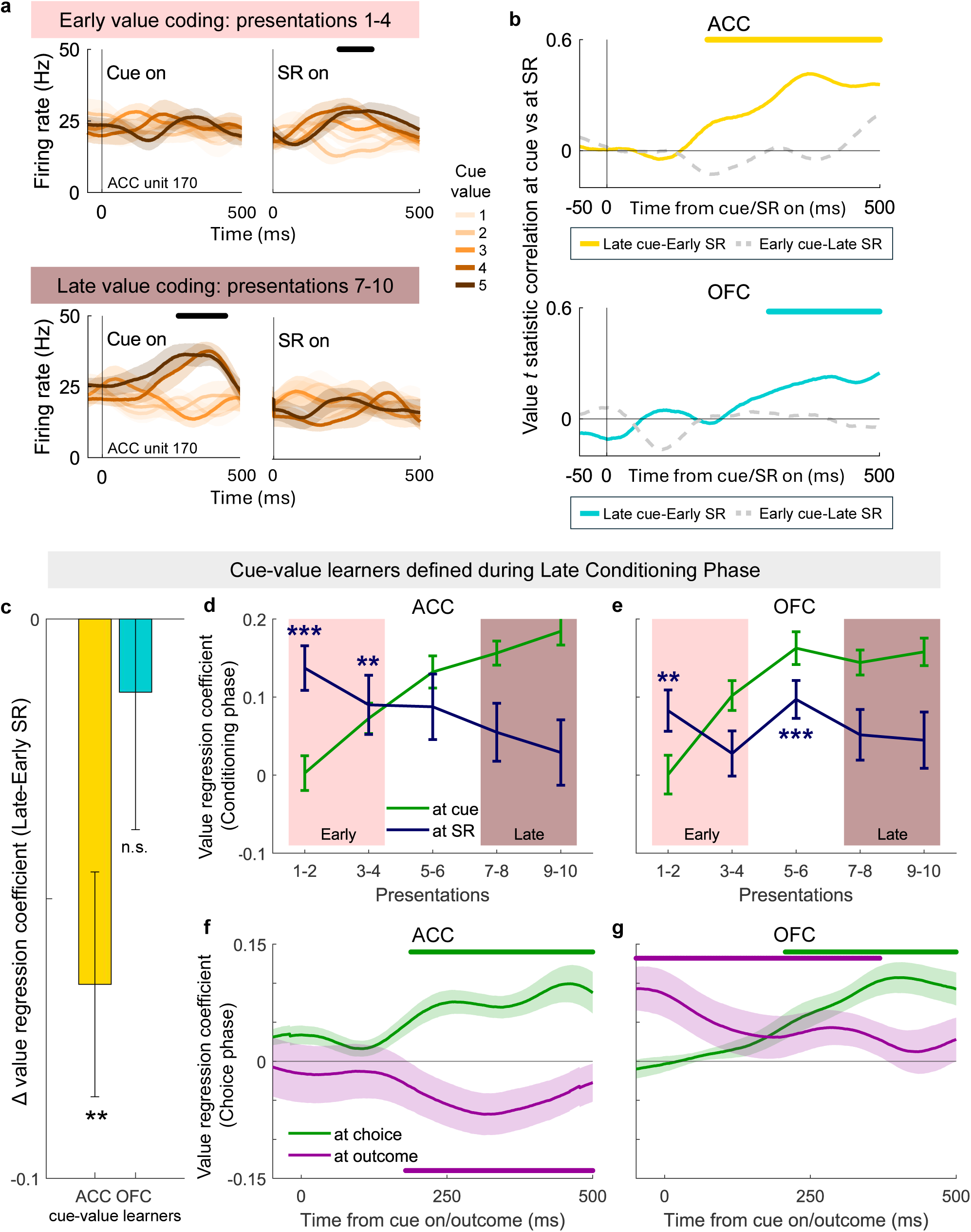
ACC neurons demonstrate RPE-driven learning dynamics. **a**, Example ACC single unit showing value-related activity following the secondary reinforcer in the early Conditioning trials (top, cue presentations 1-4); value-related activity then emerges during the cue-period in late Conditioning trials (bottom, cue presentations 7-10). Solid horizontal line represents significant value coding (|*t* stat|≥3.1 for 20ms). **b**, Diagonal of cross-correlation matrices reflecting the time-varying relationship between value coding in the cue presentation epoch and value coding in the secondary reinforcer and outcome epoch in Conditioning trials of the Conditioning phase, averaged across all ACC (top) and OFC (bottom) neurons. Solid lines represent the relationship of interest (Late value coding at cue to Early value coding at secondary reinforcer), whereas dashed lines represent the corresponding control relationship (i.e. the reverse relationship). Coloured horizontal lines denote significance within corresponding correlation, determined by cluster-based permutation testing. **c-g**, Analyses were restricted to neurons that significantly encoded value during the cue presentation epoch during presentation blocks 4-10 in Conditioning trials of the Conditioning phase (‘cue-value learners’). **c**, Difference in value regression coefficients between Early coding and Late coding during the secondary reinforcer epoch of Conditioning trials of the Conditioning phase. Regression coefficients were estimated on firing rate averaged 100-500ms after presentation of the secondary reinforcer. \*\**p*<0.001, one-sample *t*-test. **d-e**, Value regression coefficients estimated on firing rate averaged 100-500ms after presentation of the cue (green) and of the secondary reinforcer (blue line) (±SEM) in Conditioning trials of the Conditioning phase in ACC (d) and OFC (e) cue-value learners. Trials were grouped in five bins comprising two presentation blocks each to visualize the evolution of the signal over time. \*\*\**p*<0.0001, \*\**p*<0.001, t-test vs 0. **f-g**, Chosen value regression coefficients estimated on the firing rates during the cues presentation (i.e. choice) epoch and outcome epoch in probability trials of the Choice phase in ACC (f) and OFC (g) cue-value learners defined during the Conditioning phase. Solid horizontal lines denote significance within the corresponding epoch, determined by cluster-based permutation testing.

A second feature of RPE-based learning is that, upon learning to associate value with a novel cue, neurons’ responses to reward itself, or to already known reward-predicting stimuli, decrease, since reward is now entirely predicted by the preceding novel cue (Fig. 6a). This is widely described in dopaminergic neurons, but is yet to be demonstrated in frontal neurons. Although value coding at the secondary reinforcer remained evident across the ACC population in later presentations (Fig. 5), we next asked whether it diminished specifically in neurons that acquired cue-value representations. We thus tested whether neurons that acquired value representations at cue also decreased value responses associated with the secondary reinforcer (GLM-2). We found that ACC neurons that acquired value representations at cue within 4-10 presentations (‘cue-value learners’, *n* = 21; *t*(20) = 20.15, *p* < 0.0001, one-sample *t*-test on absolute *t* statistic for cue value after cue presentation) showed a significant decrease in value coding in the secondary reinforcer epoch between the Early and the Late presentations (difference between value regression coefficients was significantly different from 0; *t*(20) = −3.25, *p* = 0.004, one-sample *t*-test; Fig. 6c), whereas OFC cue-value learners did not (*n* = 21; *t*(20) = −0.53, *p* = 0.6, one-sample *t*-test on difference between value regression coefficients at cue and at secondary reinforcer; *t*(20) = 20.23, *p* < 0.0001, one-sample *t*-test on absolute *t* statistic for cue value after cue presentation). To visualise the time course of this shift in value coding from secondary reinforcer to cue, we grouped Conditioning trials in smaller bins, comprising two presentation blocks each (*n* = 20 trials/bin), and estimated value regression coefficients at both cue and secondary reinforcer (GLM-2; Fig. 6d,e). After 4 presentations, cue-value learners in ACC displayed no significant value responses at secondary reinforcer (value regression coefficients not significantly different from 0; *t*(20) = 4.80, *p* = 0.0001 at presentation 1-2; *t*(20) = 2.40, *p* = 0.03 at presentation 3-4; *p* > 0.05 at later presentations). This is coherent with RPE-based learning in ACC, since the value of the secondary reinforcer is now entirely predicted by the newly learned cue.

Finally, RPE theory predicts violations in the expectation of reward will be encoded after receipt (or omission) of reward. Encoding of discrepancies between expected and experienced reward may be demonstrated in neuronal activity by value being represented with opposite signs at choice and outcome. Indeed, the ACC cue-value learners defined during the Conditioning phase displayed this hallmark of RPE-coding during probability trials of the Choice phase, whereas OFC cue-value learners did not (Fig. 6f,g; GLM-2 and GLM-3). Altogether, these results suggest ACC and OFC neurons rapidly develop novel value representations through their existing value coding schemes, a dynamic which, in ACC neurons, follows the well-established tenets of RPE-based learning.

## Discussion

Value-guided learning and decision-making have been well-characterised in the neurons of overtrained non-human primates, and in human neuroimaging studies with often naïve participants. Yet little is known about the time course in which individual frontal neurons develop value representations, or whether frontal representations for overtrained and newly-valued stimuli are distinct or overlapping. We have demonstrated that value representations emerged in ACC and OFC neurons as rapidly as within 4-7 presentations of novel reward-predicting cues. Novel value signals resembled existing value representations associated with known cues, particularly strongly within ACC. In ACC neurons, their emergence was paralleled by a decrease in value responses associated with well-known cues, as predicted by RPE-learning theory and described in dopamine neurons. We showed that ACC neurons used a common value code when faced with choices between novel or between well-learned cues, whereas choice experience induced the recruitment of distinct neuronal subpopulations in OFC. Frontal neurons, particularly within OFC, further signalled choice novelty independently of value, with novelty signals emerging before chosen value representations.

Together, these findings bear directly on a central assumption in the decision-making field: that the single-neuron responses described in overtrained non-human primates provide a neural substrate for the rapidly arising value signals observed in human neuroimaging. Our study provides some evidence to bolster this assumption. Firstly, we showed value-related responses may emerge as rapidly as within 4-7 presentations of a novel cue. This timescale is congruent with that of a human participant undergoing a quick practice block of a task before heading into the scanner. One important parallel is that, in human studies, linguistic instructions often establish the relevant task set before learning begins (Castro-Rodrigues et al., 2022). In our task, the subjects’ extensive experience with the broader conditioning and choice structure may provide an analogous scaffold or state space, allowing value acquisition for new cues to proceed particularly rapidly. Consistent with this, rapid acquisition of novel cue-rule associations has previously been observed in dorsolateral PFC neurons when animals are well-trained on the task structure (Cromer et al., 2011), and dopamine neurons can similarly acquire value responses to new stimuli via secondary reinforcers over the course of repeated exposures (Hollerman & Schultz, 1998). This may distinguish rapid learning within a familiar task structure from settings in which animals must learn the task itself from scratch.

Secondly, we showed that value coding in ACC neurons was not altered as a function of whether the choice option presented had been experienced thousands of times before, or for the first time on that same day. In fact, ACC neurons were more likely to encode value during both choice types, rather than only during one, and were more likely to do so than OFC neurons. One possible explanation of ACC’s generalised value code across task-related variables, particularly at outcome, may be to consider the region’s functional proximity to the process of action selection (Cai & Padoa-Schioppa, 2021; Kennerley et al., 2006; Rushworth et al., 2011). ACC neurons are at the interface of reward processing and motor output (Klein-Flugge et al., 2022; Van Hoesen et al., 1993) and encode both reward and motor contingencies (Chen et al., 2024; Hayden & Platt, 2010), often doing so in a ‘common currency’ across different attributes (e.g. Kennerley et al., 2011). Abstracting information regarding an option’s value from contextual variables irrelevant for immediate choice (namely, level of training) may thus provide a more sensible substrate for action selection.

It should be noted that the value coding in ACC that generalises from novel to overtrained stimuli is typically *chosen value* coding, rather than *action value* coding. While chosen value coding has sometimes been posited as a ‘post-decision’ variable (Blanchard & Hayden, 2014), other interpretations suggest it is better thought of as reflecting the speed at which a decision process unfolds over time (Hunt et al., 2015). Within ACC, this dynamical process may be the ultimate decision itself (over which option to choose), or it may instead reflect internal decisions over where next to allocate attention or sample information (Banaie Boroujeni & Womelsdorf, 2023; Butler et al., in press; Kaanders et al., 2021).

OFC responses, on the other hand, present a case of modulation of PFC computations that may be concealed by standard non-human primate training protocols. For instance, OFC neurons were significantly less likely to encode value in a general fashion across different levels of experience, when compared to ACC. This differential neuronal recruitment may be difficult to detect by human neuroimaging methods such as fMRI. Furthermore, we found that OFC neurons were more likely to be selective for value during novel than overtrained choice. This is coherent with the preferential recruitment of the human medial frontal cortex during information sampling and consideration of novel experiences (Barron et al., 2013; Kaanders et al., 2021).

It should be noted that, while our subjects learned novel cues every session, they had already been trained on the task structure. This is unlike human participants, who are usually tasked with learning the task structure, as well as the option values, only shortly before entering the scanner. It will be interesting for future work to test the influence of a strong prior for task structure on the emergence of novel value representations.

These experience-dependent differences in OFC raise the possibility that novelty itself acts as a contextual signal for value computation. Indeed, we found that the training level on a given option, i.e. its novelty *per se*, modulated frontal activity independently of value, particularly in OFC. Orthogonality to value is a common feature of OFC coding for other task-related variables, such as informativeness (Blanchard et al., 2015; Bussell et al., 2024), stimulus identity (Klein-Flugge et al., 2013), choice risk (O’Neill & Schultz, 2010) or social category (Barat et al., 2018), and prefrontal novelty detection-related responses are similarly uncorrelated with reward-related activity (Zhang et al., 2022). Our temporal analysis further showed that novelty signals preceded chosen-value signals across frontal regions, suggesting that novelty may provide an early contextual signal that shapes subsequent value computation. This interpretation fits with accounts emphasizing the role of prefrontal cortex, and OFC in particular, in representing task-state structure (Boorman et al., 2021; Schuck et al., 2016; Veselic, Gutierrez, et al., 2025; Veselic, Muller, et al., 2025; Wilson et al., 2014). Elston and Wallis (2025) showed that when identical cues carried different values across task states, OFC recruited distinct neuronal subpopulations, revealing context-dependent coding. Here, novelty may similarly mark the current choice context (“new” vs “well-learned”), engaging partially separable subpopulations to encode value conditional on that context. Although novelty was not strictly relevant for correct choice once values were learned, it may still index uncertainty about cue-outcome mappings or value estimates, thereby supporting the allocation of attention and information gathering when uncertainty is high (Hunt et al., 2018).

Beyond showing that novelty contextualises frontal value coding, our results also speak to the mechanism by which these representations are acquired. We report that RPE-based learning in primate ACC neurons follows the same pattern as that observed in dopaminergic RPE neurons. Whereas RPE-coding during learning of stimuli and action values has been previously shown in ACC neurons (Kennerley et al., 2011; Matsumoto, Matsumoto, et al., 2007; Muller et al., 2024; Seo & Lee, 2007), here we specifically characterise cue-value learning as a temporal shift in value responses from known to newly learned reward-predicting cues (Amo et al., 2022; Sutton, 1988). We further showed that ACC neurons can acquire novel value representations as rapidly as within 4-7 trials, a timescale that may enable within-session – or, in fact, within-block – behavioural adaptation (Cole et al., 2024). This aligns with broader primate accounts of ACC integrating learning signals across timescales and uncertainty (Cavanagh et al., 2016; Kennerley et al., 2011; Monosov et al., 2020), and with recent evidence that ACC value signals are not purely reward codes, but may reflect structural task features (Miranda et al., 2026). Coupled with the notion that ACC is a primary cortical recipient of dopaminergic input (Williams & Goldman-Rakic, 1998), such rapid RPE-like learning may allow for our findings to be interpreted through the lens of meta-reinforcement learning, or Meta-RL (Wang et al., 2018). Meta-RL implements a slow, dopamine-driven RL algorithm to change the weights in a recurrent neural network which learns to implement a second, fast, PFC-based RL algorithm that changes trial-to-trial in the activation of the units. Within the context of our task, this may translate to dopaminergic input acting over several sessions to learn the structure of the task (i.e. “learning novel cues during conditioning will maximise reward during choice”), while PFC learning occurs rapidly within-session to establish cue-value associations and support behavioural adaptation (Hattori et al., 2023). While our study provides evidence in support of PFC’s rapid learning timescale, future work ought to parse the relative dopaminergic and cortical contributions to RPE-based learning by selectively interfering with the different circuit components. Furthermore, ACC may support richer forms of RPE-based learning than a single mean value update: recent work suggests that ACC neurons differ in their relative sensitivity to positive and negative prediction errors, producing a spectrum of value estimates that can collectively encode the reward distribution (Muller et al., 2024). Investigating how reward distributions are rapidly learned will be an interesting avenue for future research.

In summary, our findings show that frontal value signals can emerge within only a few experiences with a novel option to support newly encountered choices. ACC exhibited a generalised value code with RPE-like learning dynamics, whereas OFC’s neural code was more sensitive to novelty and choice context. These results support the translational relevance of primate single-neuron studies of value-guided behaviour, and reveal regional differences that may be difficult to detect with neuroimaging alone. They further highlight the need for closer alignment of behavioural paradigms across species to build a more unified account of FC contributions to learning and decision-making.

## Acknowledgements

We would like to thank Sean Cavanagh and Viktor Kewenig for earlier contributions.

## Author contributions

Conceptualization: L.T.H., W.M.N.M., S.W.K. Methodology: L.T.H., W.M.N.M., S.W.K. Investigation: L.T.H., W.M.N.M., S.W.K., E.G., T.H.M., J.L.B., S.V. Visualization: E.G. Funding acquisition: S.W.K. Project administration: S.W.K. Supervision: S.V., S.W.K. Writing – original draft: E.G. Writing – review & editing: T.H.M., J.L.B., L.T.H., S.V., S.W.K.

## Data and materials availability

All data and code to reproduce figures will be available at https://github.com/elenagutierr/learning_paper upon publication.

## Methods

### Subjects

Two adult male rhesus macaques (*M. mulatta*), M and F, were used as subjects in the study. They weighed 7-10 kg at the time of neuronal data collection and were ∼4 years old at the start of the experiment. Their daily fluid intake was regulated to maintain subject motivation. All experimental procedures were approved by the UCL Local Ethical Procedures Committee and the UK Home Office (PPL Number 70/7231) and carried out in accordance with the UK Animals (Scientific Procedures) Act.

### Behavioural protocol

The details of the experimental set up have been described previously (Hunt et al., 2018). The task was delivered through the MATLAB-based toolbox MonkeyLogic (http://www.monkeylogic.net/, Brown University) (Asaad & Eskandar, 2008a, 2008b; Asaad et al., 2013). Eye position was monitored using an infrared camera (ISCAN ETL-200) sampled at 240 Hz. Subject responses during the task were made with a voltage-gating joystick (APEM). Joystick and eye positions were relayed to MonkeyLogic for use online during the task, and were also recorded by MonkeyLogic at 1,000 Hz for offline analysis. Juice was delivered to a spout placed to the subjects’ lips through a peristaltic pump (ISMATEC IPC).

### Task

The details of the task have been described previously (Cavanagh et al., 2019). During each session, subjects underwent a short conditioning procedure (Conditioning phase) and then performed a value-based decision-making task (Choice phase). During the Conditioning phase they were taught the value of ten Novel cues, five associated with receiving different quantities of reward (magnitude cues, predicting 0.14g, 0.33g, 0.51g, 0.71g or 0.90g of juice) and five associated with different probabilities of receiving a fixed amount of reward (probability cues, predicting 10%, 30%, 50%, 70% or 90% probability of 0.51g of juice). The conditioning procedure consisted of twenty blocks of one-alternative ‘forced choice’ trials (Conditioning trials), alternating between magnitude and probability cues, each followed by two binary choice trials between two cues of the attribute at hand. Within a single block, each of the value levels of the given attribute was presented once. Ten blocks were completed for each attribute.

The Conditioning phase was followed by a Choice phase, where subjects were presented two-alternative choices. Choice options included the Novel cues learned that day during the Conditioning Phase, as well as ten Overtrained (OT) cues (five predicting magnitude of reward and five predicting probability of reward). These were stimuli which the subjects had been heavily exposed to in training sessions prior to data collection (M: ∼1500, F: ∼3000 exposures to the stimulus set). The same OT stimulus set was used for every recording session, whereas Novel cues were exclusively used for one session. Choice trials comprised choices between two OT cues (OT trials), two Novel cues (Novel trials) or one cue from each stimulus set (Mixed trials). Choices were only offered within attribute and never between cues of equal value. Choice accuracy was therefore defined as selection of the higher-value cue. All trial types were pseudo-randomly interleaved.

All trials were initiated by returning the joystick to centre position. During a Conditioning trial, subjects were first required to fixate a central red fixation point (size: 0.5 × 0.5 visual degrees) amid a white background for 500 ms (fixation radius: 3 visual degrees). If central fixation was not acquired within a 10s window a short ‘timeout’ period was given before the trial restarted. Upon successful fixation, the fixation point disappeared and a single iso-luminant picture cue (100 x 100 pixels) was presented 6.5 visual degrees to the left or the right of the centre of screen. Choice was required through leftward or rightward joystick movement within a 5s window, otherwise the trial was aborted. Upon choice of the cue, this was highlighted by drawing a grey square around it for 500ms. A grey square thereby replaced the cue for a further 500ms. This was followed by presentation for 500 ms of a secondary reinforcer, which was a coloured bar whose height indicated the value associated with the chosen stimulus. A further 500ms pre-feedback period was then followed by the reward epoch, which included reward delivery via a juice pump.

During Choice trials, following central fixation, two cues appeared to the right and the left of the screen. Subjects were free to saccade anywhere on or off the screen and choose one of the cues through joystick movement. Once a response was acquired, the chosen cue was highlighted by drawing a grey square around it for a 500ms pre-feedback period. The unchosen cue thereby disappeared from the screen, and the reward epoch (including juice delivery) was initiated.

After completion of the present task on ‘Day 1’, subjects went on to perform a different decision-making task on the subsequent three sessions (‘Day 2-4’) – described by Hunt et al. (2018) – which used the Novel cues learned on Day 1. The session schedule thereby restarted with a new Day 1 session. Here we only report on data collected on Day 1 sessions.

### Neuronal recordings

Details of our methods for neurophysiological recordings were reported elsewhere (Hunt et al., 2018). Here we report on data recorded from two target regions: ACC and OFC. We defined ACC as the entire dorsal bank of the anterior cingulate sulcus from AP 27-37 mm. All neurons lateral to the medial orbital sulcus and medial to the lateral orbital sulcus were considered OFC.

### Data analysis

#### Behavioural analysis

Behavioural data for the Choice phase have been reported previously (Cavanagh et al., 2019). All analyses collapsed trials across attributes, unless otherwise specified. We assessed increase in choice accuracy during the Conditioning phase, as proxy for learning of the Novel cues, by testing whether accuracy was significantly above chance in the Conditioning Phase choice trials (pooled across attributes) within bins of two blocks/bin.

#### Neuronal analyses

For all neuronal analyses, firing rates were smoothed with a Gaussian smoothing window of 250ms (50ms standard deviation) and normalised. Analyses using a boxcar method for smoothing (averaging firing rates within a sliding 200ms window in 10ms steps) produced comparable results.

#### General linear models

We used three different GLMs, detailed in Table 1. GLM-1 included Novel and OT trials of the Choice phase and examined how neurons’ selectivity to value was influenced by the level of training on a set of cues (Fig. 3). The same GLM was also used to test whether neurons were selective to the novelty of a choice, independently of the value of the cues presented (Fig. 4). GLM-2 modelled the relationship between the neurons’ firing rate and the value of the cues presented (Fig. 5-6). We fit the same GLM to three splits of the conditioning trials across the Conditioning Phase, which included cue presentation blocks 1-4, 4-7 and 7-10 respectively (*n*=40 trials/bin) (Fig. 5, 6a-b). We also separately fit GLM-2 to Conditioning trials from cue presentation blocks 4-10 in order to define neurons that acquired cue-related value selectivity during the Conditioning phase (‘cue-value learners’, Fig. 6c-e). In the same neurons, we also tested the presence of opponent coding for value during the Choice phase by fitting GLM-2 to firing rates during the cue presentation epoch, and GLM-3 to firing rates during the outcome epoch (Fig. 6f,g; see *Reward prediction error-coding* below for details). For GLM-3, interaction terms were orthogonalised with respect to their lower-order terms to remove shared variance; that is, probability of reward on rewarded and unrewarded trials were orthogonalised relative to categorical reward delivery.

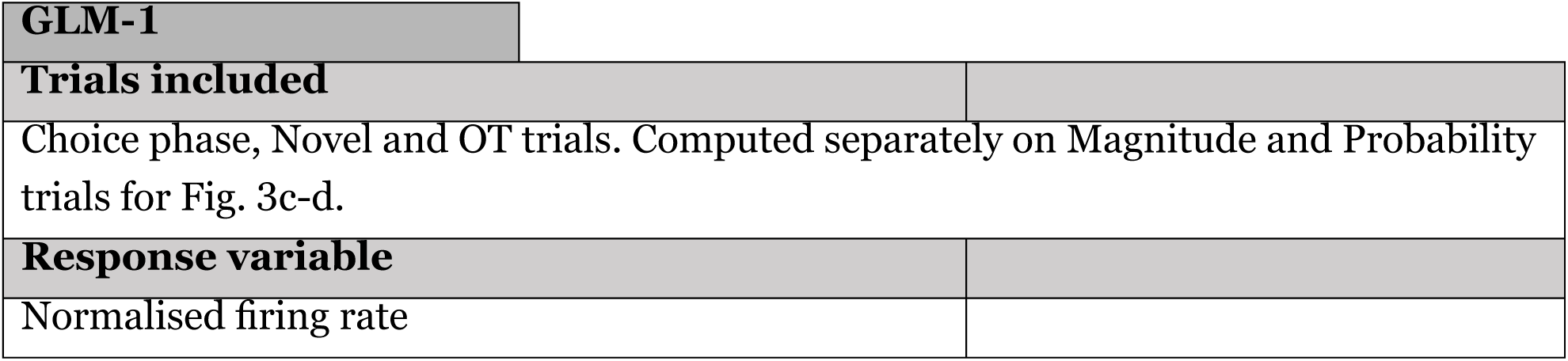
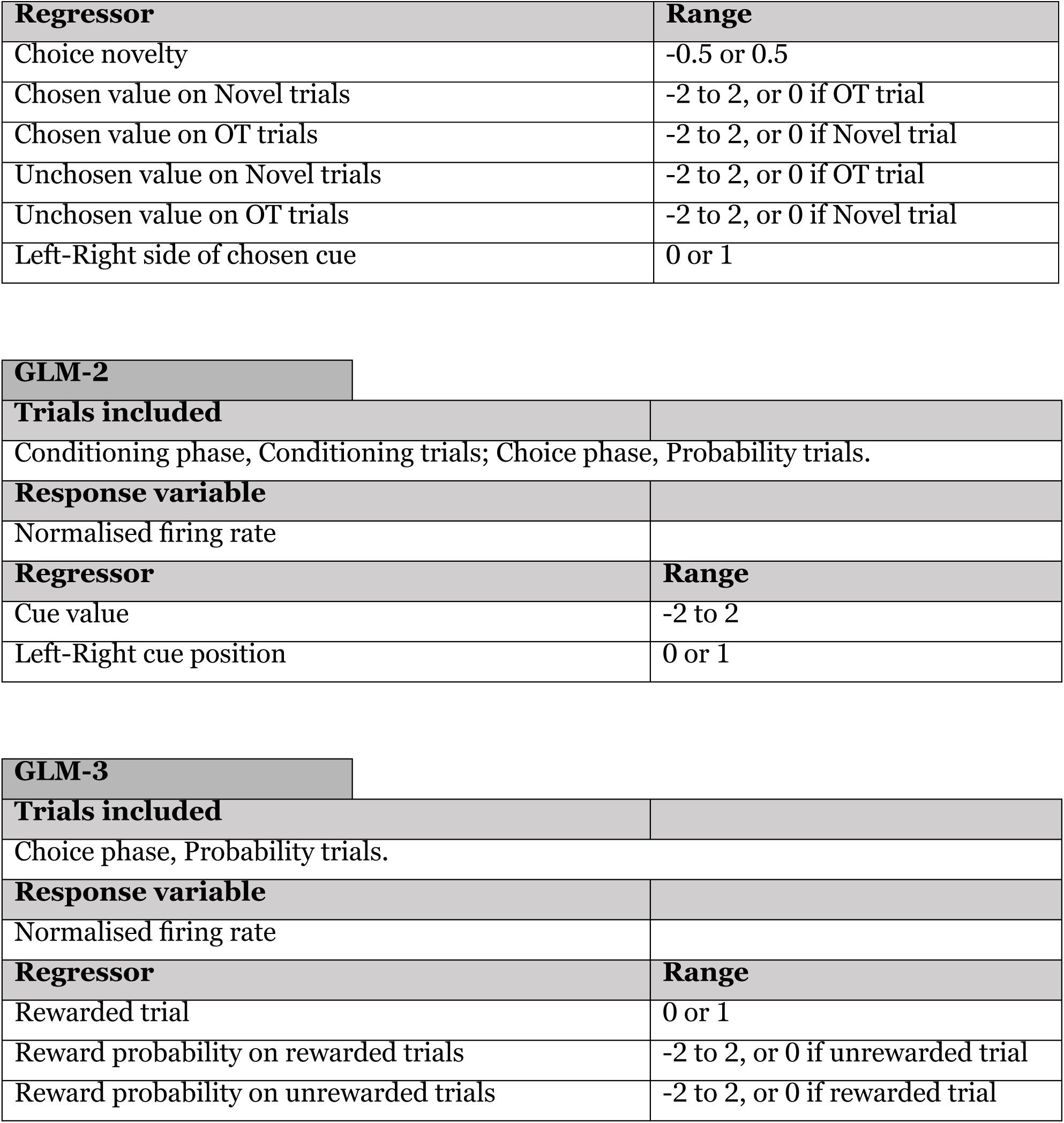

| GLM-1 |
| --- |
| Trials included |
| Choice phase, Novel and OT trials. Computed separately on Magnitude and Probability trials for Fig. 3c-d. |
| Response variable |
| Normalised firing rate |

| **Regressor** | **Range** |
| --- | --- |
| Choice novelty | -0.5 or 0.5 |
| Chosen value on Novel trials | -2 to 2, or 0 if OT trial |
| Chosen value on OT trials | -2 to 2, or 0 if Novel trial |
| Unchosen value on Novel trials | -2 to 2, or 0 if OT trial |
| Unchosen value on OT trials | -2 to 2, or 0 if Novel trial |
| Left-Right side of chosen cue | 0 or 1 |

| **GLM-2** |  |
| --- | --- |
| **Trials included** |  |
| Conditioning phase, Conditioning trials; Choice phase, Probability trials. |  |
| **Response variable** |  |
| Normalised firing rate |  |
| **Regressor** | **Range** |
| Cue value | -2 to 2 |
| Left-Right cue position | 0 or 1 |

| **GLM-3** |  |
| --- | --- |
| **Trials included** |  |
| Choice phase, Probability trials. |  |
| **Response variable** |  |
| Normalised firing rate |  |
| **Regressor** | **Range** |
| Rewarded trial | 0 or 1 |
| Reward probability on rewarded trials | -2 to 2, or 0 if unrewarded trial |
| Reward probability on unrewarded trials | -2 to 2, or 0 if rewarded trial |

*Cross-correlation of parameter estimates from GLMs (Fig. 3b,d, Fig. 6b).* After the models above were estimated for each neuron, we correlated the *t* statistics associated with parameter estimates for different regressors across neurons. In Fig. 3b and Fig. 6b, parameter estimates were taken from 50ms pre-stimulus presentation (i.e. presentation of the cue(s) or of secondary reinforcer) or event (i.e. response in the choice epoch or reward delivery in the outcome epoch) to 500ms post-stimulus presentation or event, and correlations were repeated on all possible combinations of timepoints to produce a cross-correlation matrix. In the case of Fig. 6b, only the diagonal of the cross-correlation matrix was included for further analysis for clarity purposes. In Fig. 3d, firing rates were averaged in the 1.5s following cue onset (i.e. the same post-cue choice epoch shown in Fig. 3b), and correlations were estimated on the resulting parameter estimates. Additionally, parameters were estimated separately between attributes by fitting GLM-1 separately to firing rate on Magnitude and Probability trials.

In Fig. 3, the *t* statistics for the chosen value on Novel trials and chosen value on OT trials regressors from GLM-1 were correlated to assess similarity between neural representations associated with differing levels of training.

In Fig. 6, the *t* statistics for the cue value regressor from GLM-2 were used to assess similarity between emerging neuronal representations during the cue presentation epoch and existing representations during the secondary reinforcer presentation epoch. We correlated the value *t* statistics obtained by fitting GLM-2 to firing rates from trials within presentation blocks 7-10 during the cue presentation epoch to the *t* statistics obtained by fitting GLM-2 to firing rates from trials within blocks 1-4 during the secondary reinforcer epoch. We further compared the data to the opposite correlation as a control – between *t* statistics obtained by fitting GLM-2 to firing rates from trials within presentation blocks 1-4 during the cue presentation epoch to the *t* statistics obtained by fitting GLM-2 to firing rates from trials within blocks 7-10 during the secondary reinforcer epoch. This procedure allowed us to ascertain that the positive correlation in the data was only present between the hypothesised presentation blocks.

*Statistical inference on cross-correlation of parameter estimates (Fig. 3b, Fig. 6b).* We used a cluster-based permutation test approach to assess whether high/low correlation coefficients were significantly greater than expected by chance (Hunt et al., 2018). For Fig. 3b and 6b, we permuted the regressors of GLM-1 and GLM-2 (1,000 permutations). A null distribution of correlation coefficients was obtained by storing the 97.5^th^ percentile at each timepoint of the permuted data. For each permutation, we identified clusters that exceeded the null distribution and the size of the largest cluster was stored. We used the 97.5^th^ percentile of this distribution as a threshold for deeming whether the clusters in the data exceeding the null distribution were significant. In Fig. 6b, a significant difference between the positive correlation in the data and the control was also assessed through the same procedure, but where the null distribution was obtained by permuting the labels of the presentation blocks (i.e. 1-4 and 7-10) once the parameters had been estimated and correlated. In Fig. 3b, we also tested a significant difference in correlation coefficients between ACC and OFC neurons by permuting the labels of the regions once parameters were estimated (Supplementary Fig. 2).

*Estimating temporal structure of choice novelty and chosen value codes (Fig. 4d).* To assess whether the choice novelty code temporally preceded the chosen value code, we quantified the lagged relationship between choice novelty and chosen value CPD estimates. For each neuron, choice novelty and chosen value CPD time courses during Novel trials (−500ms to +1s from choice onset) were extracted using GLM-1 and z-scored across time. This was done to prevent differences in overall CPD magnitude across neurons from dominating cross-correlation estimates. We then computed the normalized cross-correlation between the two CPD time courses at lags up to 1500 ms in both temporal directions. This produced, for each neuron, a cross-correlation function describing the similarity between novelty and chosen value coding at different temporal offsets.

Because CPD estimates are unsigned, this analysis tests whether the strength of novelty coding and chosen value coding covaried across time, rather than whether the two variables were encoded with the same sign. We therefore quantified temporal asymmetry by comparing cross-correlations at matched positive and negative lags. Positive forward-backward asymmetry values indicate stronger alignment when the choice novelty code precedes the chosen value code, whereas negative values indicate stronger alignment when chosen value coding precedes choice novelty coding.

Statistical significance was assessed with a permutation test in which novelty CPD time courses were randomly reassigned across neurons relative to chosen value CPD time courses. For each of 500 permutations, we recomputed the forward-backward asymmetry and averaged it across neurons. The significance threshold was defined as the maximum, across lags, of the 95th percentile of this null distribution, thereby correcting for multiple lag comparisons. Observed lags exceeding this threshold were considered significant.

*Reward prediction error-coding (Fig. 6c-g).* We restricted these analyses to neurons selective for value during conditioning trials within the cue presentation blocks 4-10 of the Conditioning phase (‘cue-value learners’). Our approach to assess the presence of RPE-coding at the population level was two-fold. In Fig. 6c, we fit GLM-2 to firing rates from conditioning trials within blocks 1-4 and 7-10, averaged 100-500ms after secondary reinforcer presentation, and tested whether the difference between value regression coefficients averaged across the population was significantly above or below zero (one-sample *t*-test, *p* < 0.05). In Fig. 6f,g, we fit GLM-3 to firing rates from probability trials of the Choice phase during the outcome epoch. In order to determine whether the value regression coefficients averaged across the population were greater/lower than expected by chance, we used the same cluster-based permutation test described above, where the null distribution was obtained by permuting the regressors. To account for the prefrontal and cingulate heterogeneity of positive and negative value coders, the sign of the value regression coefficient of interest was flipped for neurons that displayed a negative value regression coefficient (GLM-2) in a control epoch. In Fig. 6c, control value regression coefficients were determined during the cue presentation epoch of conditioning trials of the Conditioning phase. In Fig. 6f,g, control value regression coefficients were determined during the cue presentation epoch of probability trials of the Choice phase.

*Neuronal selectivity (Fig. 3e, 4b, 6c-g).* Neuronal selectivity was quantified by fitting a GLM to each neuron’s firing rate averaged over the analysis window of interest. For whole-trial analyses (Figs. 3-4), this was the full 1.5s period following cue onset. For event-specific analyses (Fig. 6), firing rates were averaged in 100-500 ms windows following cue onset or secondary reinforcer presentation. A neuron was considered selective for a given parameter at *p* < 0.05 for the corresponding parameter estimate. Binomial tests with a false positive rate of *p* < 0.05 were used to determine whether a region comprised a significant proportion of neurons selective for a given parameter (or a false positive rate of *p* < 0.0025 to determine selectivity for two parameters). When testing if a region comprised a significant proportion of neurons selective for a given parameter at two different epochs, the false positive rate was determined within-region by computing false positive rates using permuted data. Proportions of selective neurons were compared using a chi-squared test.

For visualisation purposes (Fig. 4c and Fig. 6a) we also estimated neuronal selectivity over time. To do this, we fit a GLM to each neuron’s firing rate at each time-point. We defined significant selectivity at a given timepoint as an absolute *t* statistic≥3.1 for 20 consecutive time-points (Kennerley et al., 2011). This criterion produced acceptable type I (false positive) error levels when examining the proportion of neurons reaching criterion during the fixation epoch (500ms window pre-cue presentation), with less than 2% neurons crossing the criterion by chance.

*CPD analysis and statistical inference (Fig. 3a, Fig. 4a, Fig. 5)*. The contribution of the regressors in the three GLMs to explaining variance in neurons’ firing rates was assessed by calculating the CPD for each regressor over time, which is defined for explanatory variable *X_i_* as follows:

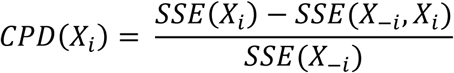

where *SSE*(*X*) refers to the sum of squared errors in a general linear model (GLM) that includes a set of EVs *X*, and *X*_−*i*_ is a set of all the EVs included in the full model except for *X_i_* (Cai et al., 2011; Kennerley et al., 2011). CPD was assessed to be higher than expected by chance through the same permutation test procedure described above. When determining statistical significance within a given CPD trace, for each of 1,000 permutations, the GLMs’ parameters were permuted when estimating CPD. A null distribution of population CPDs was obtained by storing the 95^th^ percentile at each timepoint of the permuted data across neurons. For each permutation, the maximum length of consecutive timepoints exceeding the null distribution was stored. We used the 95^th^ percentile of this distribution as a threshold for deeming whether the cluster sizes in the data were significant. To determine a significant difference between different CPD traces, the null distribution was obtained by permuting the labels of the data once the parameters had been estimated: in Fig. 3a, it was the Novel trials or OT trials label; in Fig. 4a it was the label of the region (ACC or OFC); in Fig. 5, this was the presentation block 1-4 or 7-10 label.

