## Supplemental Materials for "Rapid value learning reveals generalized and context-dependent codes in frontal cortex"

### Supplementary materials

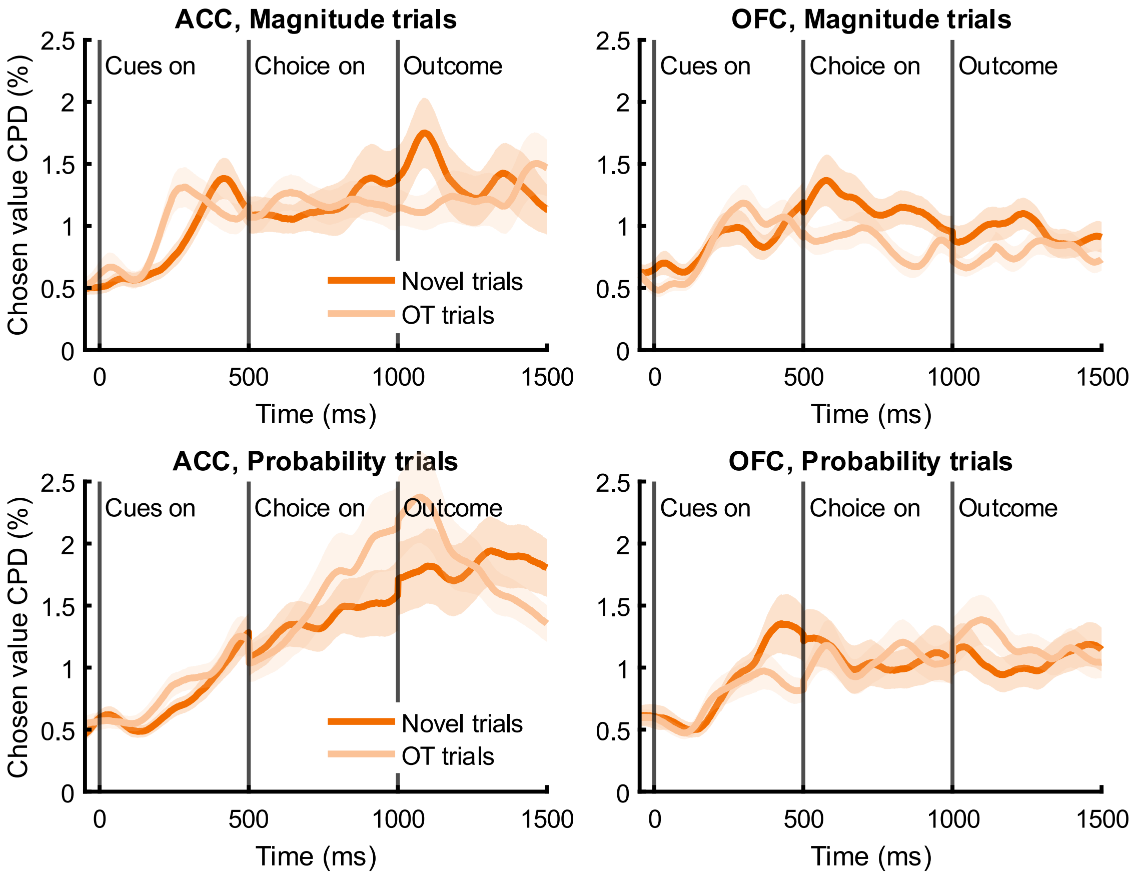


**Supplementary Figure 1. Chosen value coding within attributes during Novel and OT trials.** Same analyses as Figure 3a, but split according to reward attribute predicted by the cues presented, magnitude (top) or probability (bottom) across all ACC (left) and OFC (right) neurons.

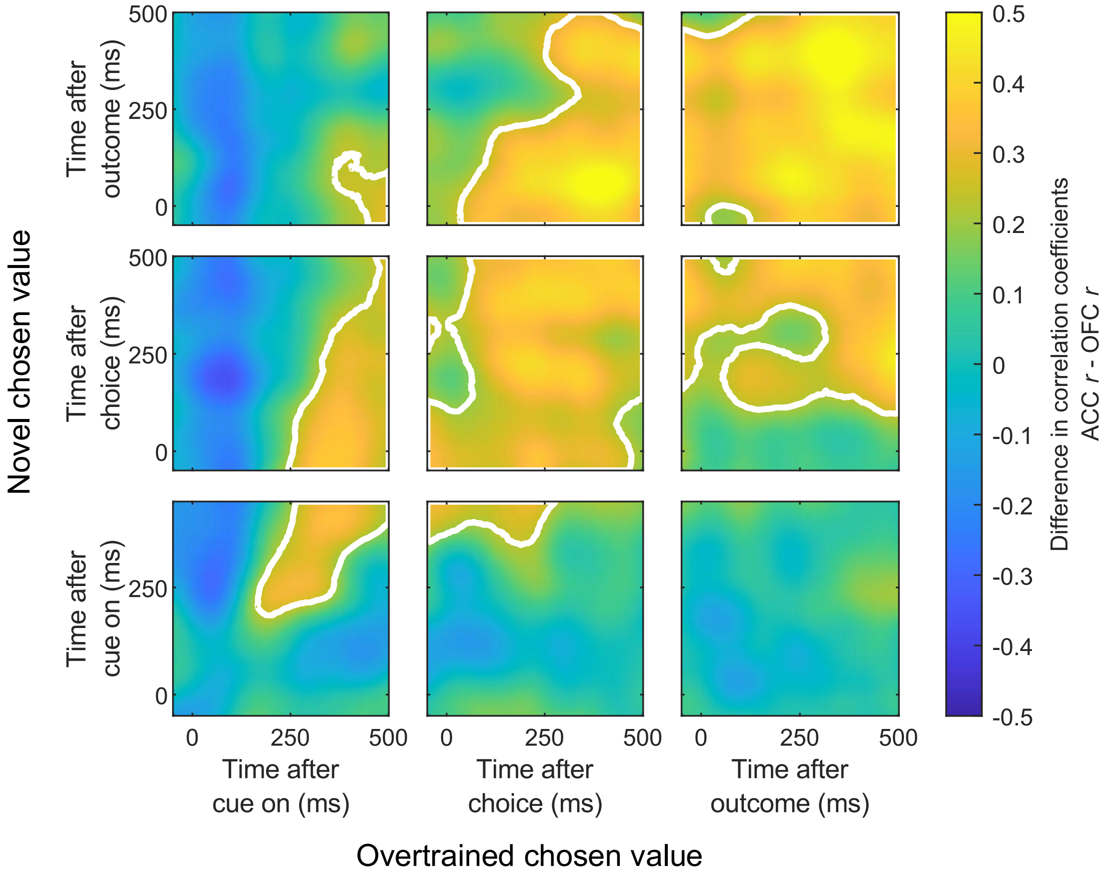


**Supplementary Figure 2.** Difference between ACC and OFC correlation coefficients (shown in Fig. 3b). White lines denote significance determined by cluster-based permutation testing.

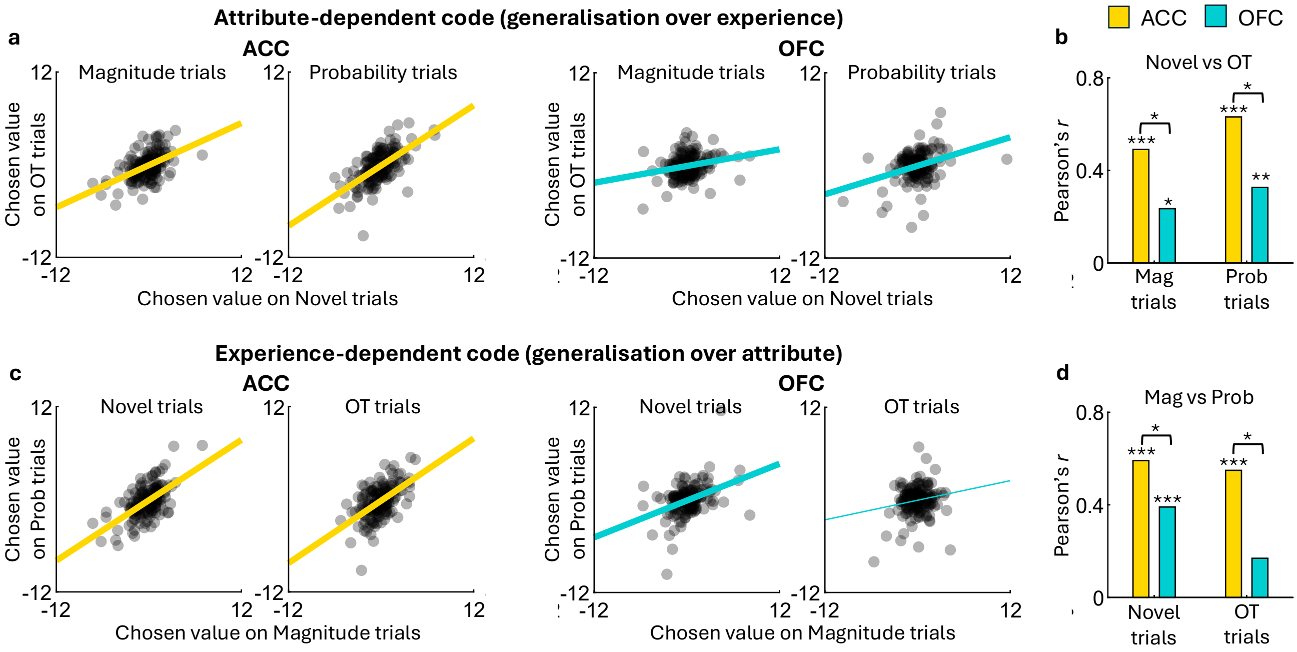


**Supplementary Figure 3. Generalisation of value coding across experience and reward attribute. a**, Neuron-wise scatter plots of chosen value *t* statistics on Novel versus OT trials, shown separately for magnitude and probability trials in ACC and OFC. Each point represents one neuron. Lines indicate least-squares fits, with thickness indicating a significant correlation. **b**, Pearson correlation coefficients for the relationships shown in a. Correlation coefficients in panel a were compared across regions using permutation tests with shuffled ACC/OFC labels ($n$=10,000). *$p < 0.05$, **$p < 0.001$, ***$p < 0.0001$. **c**, Same as panel a, but for value *t* statistics on magnitude versus probability trials, shown separately for Novel and OT trials. **d**, Same as panel b, but for the relationships shown in c.

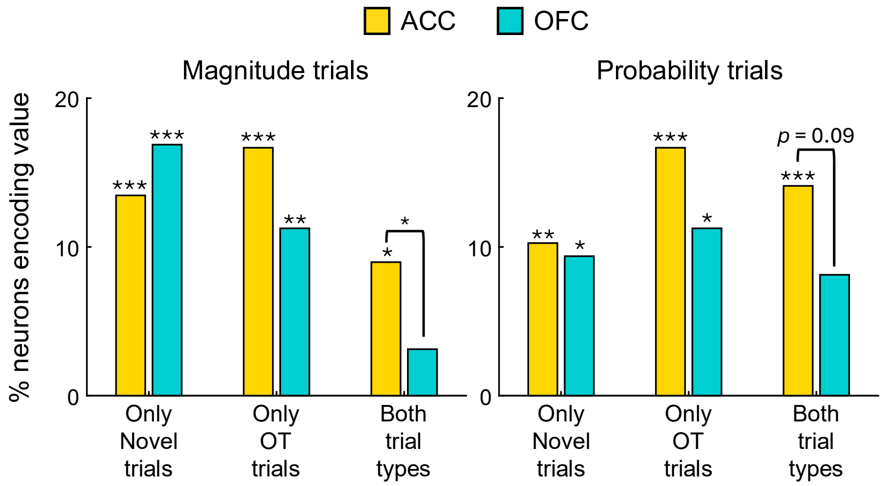


**Supplementary Figure 4.** Proportion of ACC and OFC neurons encoding value only in Novel trials, only in OT trials or in both trial types, split according to the reward attribute predicted by the cues presented, magnitude (left) or probability (right). *$p$<0.05, **$p$ < 0.001, ***$p$ < 0.0001, Binomial test and Chi-squared test, Bonferroni-corrected.

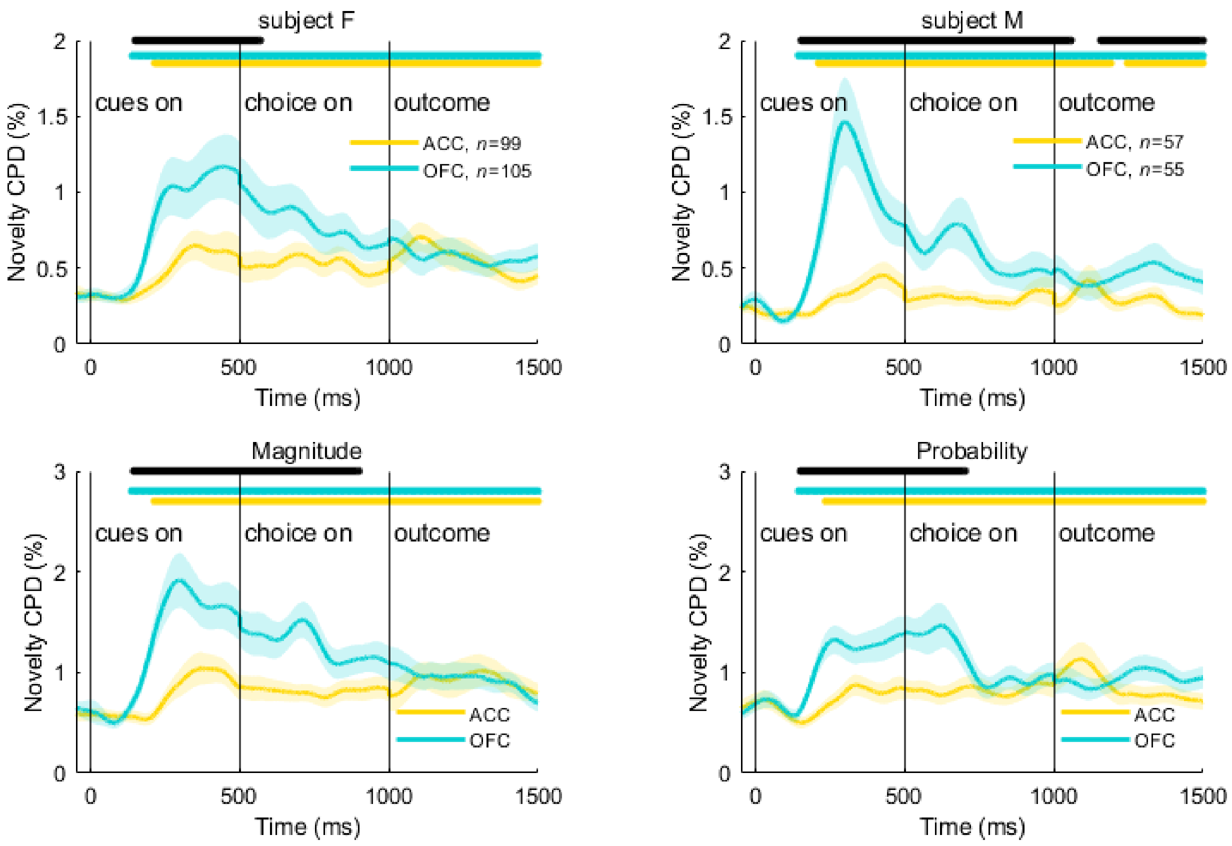


**Supplementary Figure 5. Choice novelty coding within subjects and attributes.** Same analyses as Figure 4a, but split according to subject, F (top left) or M (top right), or reward attribute predicted by the cues presented, magnitude (bottom left) or probability (bottom right).

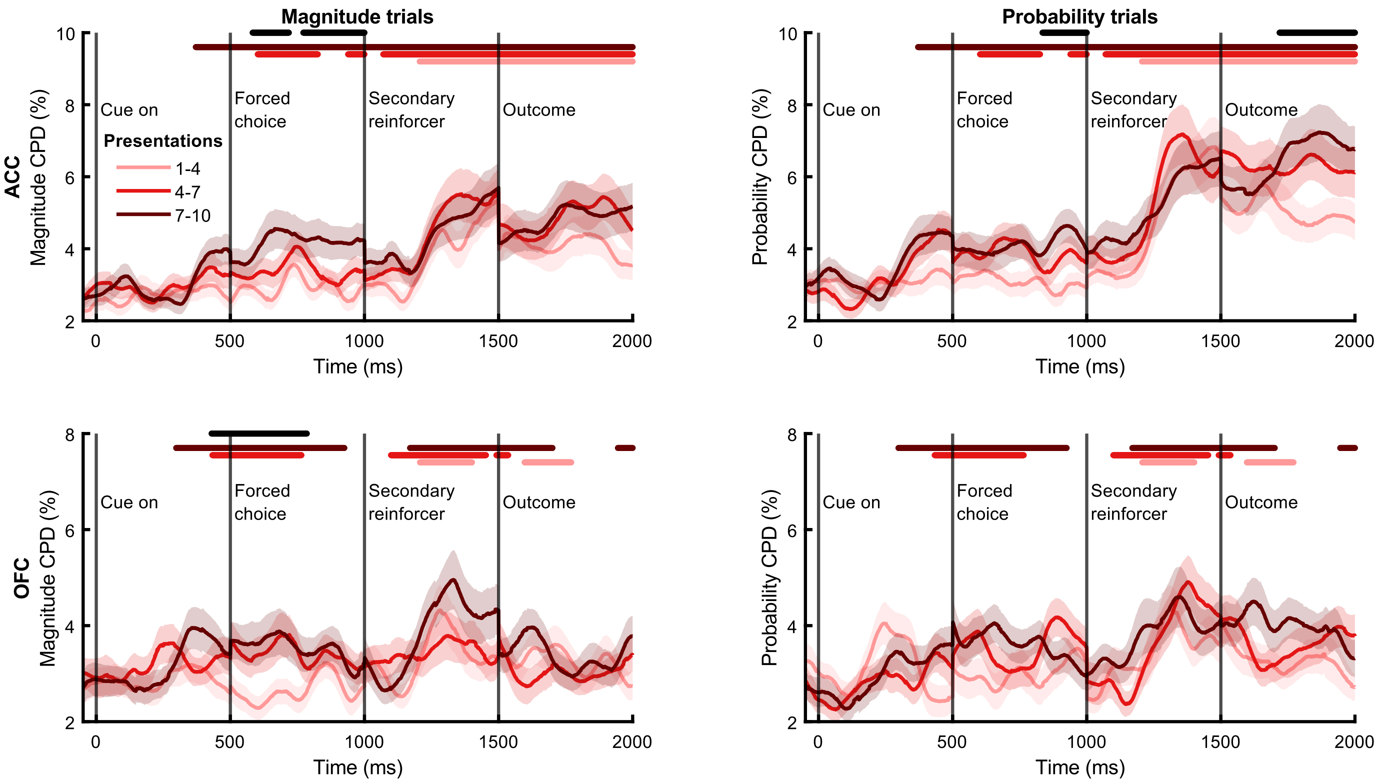


**Supplementary Figure 6. Emergence of cue-related value representations within attribute.** Same analyses as Figure 5, but trials were split according to attribute predicted by the cue presented, magnitude (left) or probability (right).
